# epiTCR-KDA: Knowledge Distillation model on Dihedral Angles for TCR-peptide prediction

**DOI:** 10.1101/2024.05.18.594806

**Authors:** My-Diem Nguyen Pham, Chinh Tran-To Su, Thanh-Nhan Nguyen, Hoai-Nghia Nguyen, Dinh Duy An Nguyen, Hoa Giang, Dinh-Thuc Nguyen, Minh-Duy Phan, Vy Nguyen

## Abstract

**Motivation:** Antigen recognition by T-cell receptors (TCRs) triggers cascades of immune responses. Successful predictions of the TCR and antigen (as peptide) bindings therefore signify the advancements in immunotherapy. However, most of current TCR-peptide interaction predictors fail to predict unseen data. This limitation may be derived from the conventional usage of TCR and/or peptide sequences as input, which may not adequately reflect their structural characteristics. Therefore, incorporating the TCR and peptide structural information into the prediction model to improve the generalizability is necessary.

**Results:** We presented epiTCR-KDA as a new predictor of TCR-peptide binding that utilises structural information, specifically the dihedral angles between the residues of both the peptide and the TCR. This structural descriptor was integrated into a model constructed using knowledge distillation to enhance its generalizability. The epiTCR-KDA demonstrated competitive prediction performance, with an AUC of 0.99 for seen data and AUC of 0.86 for unseen data. Across multiple public datasets, epiTCR-KDA consistently outperformed other predictors, such as epiTCR, NetTCR, BERTrand, TEIM-Seq, TEINet, and ImRex, maintaining a median AUC of 0.9 (ranging from 0.82 to 0.91). Further analysis of epiTCR-KDA performance indicated that the cosine similarity of the dihedral angle vectors between the unseen testing data and training data is crucial for its stable performance. In conclusion, our epiTCR-KDA model, with its capacity to predict for unseen data, has brought us one step closer toward the development of a highly effective pipeline for affordable antigen-based immunotherapy.

**Availability and implementation:** epiTCR-KDA is available on GitHub (https://github.com/ddiem-ri-4D/epiTCR-KDA)

## Introduction

The interaction between T cell receptor (TCR) and an antigenic peptide presented by human leukocyte antigen (HLA) molecules plays a pivotal role in activating the adaptive immune system. Therefore, the ability to predict the binding between a TCR and a peptide is crucial for the identification of peptides used in immunotherapy. Multiple attempts have been made to create prediction tools for TCR-peptide binding using diverse computational approaches. There are simple models such as Bayesian non-parametric model[1], Random Forest (TCRex[2], epiTCR[3]), and clustering-based models (TCRdist[4, 5]). More complex models[5, 6, 7, 8] are also proposed for the classification task. Many deep learning models (NetTCR[9], DeepTCR[7], ImRex[8], tcrpred[10]) rely on convolutional neural networks to learn the TCR and peptide patterns in each interaction. TITAN[11], ATM-TCR[12], and tcrpred[10] further evaluate the pairwise interactions by crossing the TCR and peptide patterns in an attention structure. Some other tools, particularly ERGO-I[13] and pMTnet[14], use long short-term memory (LSTM) to learn the sequential information of TCR and peptides, and autoencoder layers to simultaneously improve the data understanding and reduce the feature space. Also extracting sequence information, BERTrand[15], a language model-based model, learns the amino acid position and composition in the TCR and peptide sequences contributing to the binding. Despite many machine learning and deep learning algorithms that have been applied to predict the interactions between TCR and peptides, predicting the TCR-peptide binding is still a challenge, especially when applying to unseen data where either the sequences of TCR or peptide or both are not presented in the training dataset.

Most TCR-peptide binding predictors struggle to generalise the interaction of TCR and peptide[6, 7, 14]. The first reason is the datasets used to train and testing predictive models are limited in size or diversity, particularly when it comes to the number of peptides. It was demonstrated with NetTCR that there was a positive correlation between the model performance and the size of the training dataset[9]. The available data may not represent the full spectrum of peptide variability or skewed towards certain peptide sequence patterns [16]. Furthermore, some studies[9, 17] suffered from overoptimistic classification performance when using random data split for training and testing, inadvertently resulting in data leakage due to the presence of the same peptide sequences in both training and testing data even though the CDR3β-peptide pairs are not overlapping[16]. Therefore, careful attention is needed in dataset construction and validation, with a special focus on using unseen peptides for testing, to prevent data leakage and ensure the development of robust predictive models. The second reason is the insufficient information for the models to learn from the input pair of the amino acid sequences of the TCR CDR3β region and the peptide, of which the linear sequences do not represent the spatial information of the TCR-peptide interactions. In fact, two different TCR-peptide sequence pairs can share similar spatial information and therefore can interact in the same manners [8]. This lack of spatial information might prevent models from generalising the TCR-peptide interactions, leading to low performance on unseen data.

Many protein structure prediction tools (such as AlphaFold[18], ESMFold[19], and OmegaFold[20]) transform such linear to spatial information, which might augment the learning ability of the deep learning models. The dihedrals including phi □ angle (around the backbone N-Cα bond) and psi ψ angle (around the backbone Cα-C bond) are used to represent the 3D shape of a peptide[21, 22], therefore, can serve as good features to capture the structural information of both the TCR CDR3β and peptide, guiding the models to learn the patterns of spatial interactions.

In this study, we present epiTCR-KDA to predict the TCR-peptide binding based on a knowledge distillation model (KD)[23], which learns the spatial information from dihedral angles of both the TCR CDR3β and peptides. The epiTCR-KDA was trained on a dataset of diverse TCR and peptides, with additional known non-binding peptides (wild type) sourced from public databases. The model consistently outperformed other currently available TCR-peptide binding prediction tools. Furthermore, our epiTCR-KDA also elicited the outstanding generalization ability on unseen data.

## Methods

### Data collection and generation of non-binding TCR-peptide pairs

Binding and non-binding (TCR) CDR3β-peptide pairs were collected from McPAS-TCR[24], TBAdb[25], VDJdb[26], IEDB[27], and 10X[28]. The combined dataset contains 70,083 (2.5%) binding pairs and 2,689,709 non-binding pairs that were formed by using 1,681 unique peptides and 126,841 unique CDR3β sequences. Among 1,681 unique peptides, only 7 are exclusively found in non-binding pairs, significantly lower than the number of peptides exclusively found in binding pairs (1,637 unique peptides). The small number of unique peptides in non-binding pairs highlighted the data imbalance, likely leading to bias towards the positive “binding pairs” predictions [3]. To address this issue, we augmented the data by constructing additional non-binding CDR3β-peptide pairs. We first extracted wild-type peptide sequences from TSNAdb[29], Neodb[30], and NEPdb[31], and then randomly combined them with CDR3β sequences from previously collected data. This resulted in additional 174,944 CDR3β-peptide pairs with 2,506 unique peptides for the non-binding dataset. On the other hand, we further combined 71 CDR3β sequences from TIL[32] with the extracted wild-type peptides to make additionally 132,979 non-binding CDR3β-peptide pairs. Details can be found in Figure 1A.

**Figure 1.**
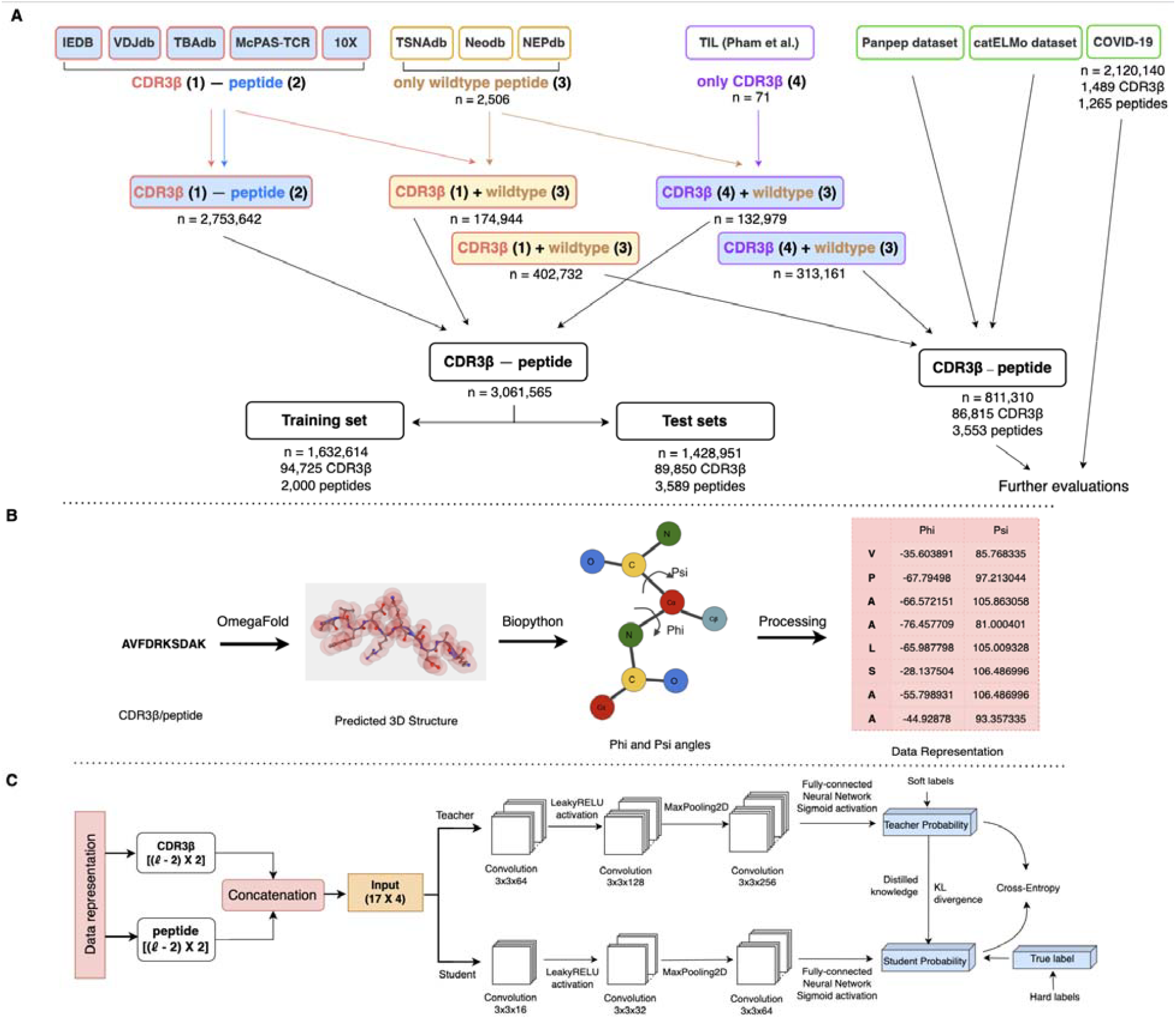
Overview of epiTCR-KDA. (A) Diagram illustrating data collection for training and evaluation of epiTCR-KDA. Five public databases (IEDB[27], VDJdb[26], TBAdb[25], McPAS-TCR[24], and 10X[28]) were collected for TCR-peptide pairs (Supplementary Table S1). Three databases (TSNAdb[29], Neodb[30], and NEPdb[31]) were gathered for self-peptides (wild-type peptides). These peptides were randomly combined with TCR from public TCR-peptide pairs to form non-binding pairs. Additionally, non-binding pairs were also generated from TIL CDR3β sequences[2] with public wildtype peptides. The data were divided into training data (Supplementary Figure S1), and testing data covering various data sources, seen and unseen peptides (Supplementary Table S2). (B) Data preprocessing steps starting from the conversion of CDR3β/peptide amino acid sequences to 3D structures using OmegaFold, followed by the calculation of the phi and psi angles, and processing this information as input for the model. (C) Structure of the KD model. The CDR3β and peptide representation (phi and psi angles) were concatenated, padded, and served as input for the KD model. The KD model involved a student model learning from the information provided by the teacher model (soft loss) and ground-truth labels (hard loss). The model was trained to predict the binding or non-binding of CDR3β-peptide pairs.

### Input data representation

The phi ([) and psi (ψ) torsion angles[33] were used to represent the structural information of both the CDR3β and peptide sequences. The CDR3β/peptide sequences were first used to model their 3D-structures using OmegaFold (version v1.1.0)[20]. The phi and psi angles were then calculated using the PDBParser function implemented in the biopython package (version 1.75)[34], resulting in (*l* − 2, 2) matrices, with *l* corresponding to the length of the sequence. The first and the last amino acids in the sequence can rotate freely around the peptide backbone, therefore their phi and psi angles were excluded. The resulting phi and psi matrix was provided to the learning model as input (Figure 1B).

### Data organization for model training and testing

The training data consisted of 1,632,614 CDR3β-peptide pairs, including 94,725 unique CDR3β sequences, and 2,000 unique peptides. The testing data comprised of 1,428,951 pairs, including 89,850 unique CDR3β sequences and 3,589 unique peptides. Of the unique peptides in the testing data, 1,948 (54.2%) were seen peptides (peptides paired with other CDR3β in the training data), and 1,641 (45.8%) were unseen peptides (peptide sequences only found in the testing data, Supplementary Table S2). The testing data were randomly split into ten testing sets, allowing the benchmark of epiTCR-KDA against other predictors. A “7 unseen dominant peptides” dataset consisting of 447,398 CDR3β-peptide pairs derived from 7 unseen peptides (Supplementary Table S3) was also randomly split into 10 subsets and used to testing the models.

### Model training

The model structure followed a knowledge distillation approach[23], akin to a teacher-student relationship (Figure 1C). The input CDR3β and peptide sequences were individually represented by matrices of phi and psi angles, which were then concatenated and padded by zeros into a 17×4 matrix (where 17 rows representing the dimension obtained from the longest sequence, and 4 columns representing the phi and psi angle pairs of the CDR3β then of the peptide, respectively). Taking this matrix as input, both the student and teacher models were built based on the convolutional neural networks (CNNs) framework. The teacher model was for binary classification, started from a convolutional layer of 64 filters of size 3×3 (with the stride of (2, 2)), followed by a LeakyReLU activation (α = 0.2), and a MaxPooling2D with 2×2 filter and stride = 1). Two convolutional layers with 128 and 256 filters (with the same filter size and stride) were subsequently applied. The output from the last layer was flattened into a 1D vector, followed by a fully connected layer, and a single unit of sigmoid activation for binary classification. The student model replicates the teacher’s prediction with reduced complexity by reusing three convolutional layers of 16, 32, and 64 filters, respectively, while other layers were kept similar to those of the teacher model. The distillation involved a Distiller object containing both models. During training, the Distiller object was compiled using Adam optimizer[33], with BinaryAccuracy metric[35] for evaluation, BinaryCrossentropy loss function[36] for the student, and KLDivergence[37] for distillation loss evaluation. These parameters were resulted from the model tuning process.

## Results

### Overview of epiTCR-KDA

To construct a prediction model for TCR-peptide binding, we tackled this problem in three main angles: collection of data, encoding of data, and structure of model. As different training datasets can severely affect the performance of the prediction model (epiTCR[3]), we focused our efforts in obtaining diverse CDR3β and peptides with known binding status from multiple publicly available sources (Figure 1A). Furthermore, we generated non-binding pairs to increase the proportion of non-binding data based on the assumption that TCR does not bind to and is not activated by human wildtype peptides (ie. self-peptides) (Figure 1A). For training data, a series of training sets was generated with an increasing number of peptides and corresponding CDR3β-peptide pairs, demonstrating that the training dataset with 2,000 unique peptide sequences exhibited the best training performance (Figure S1). Therefore, a total of 1,632,614 CDR3β-peptide pairs consisting of 94,725 unique CDR3β sequences and 2,000 unique peptides were used as the training data (Figure 1A). We hypothesize that the amino acid sequences and subsequent common encoding algorithms such as one-hot encoding or blosum62 might not provide insights into the 3D structures of the two binding partners. Therefore, we used dihedral angles as input data to better reflect the structural information of the CDR3β and peptides. Finally, we used a knowledge distillation (KD) model to learn from the dihedral angle matrix input (Figure 1C). Mimicking the process of transferring knowledge from the teacher to the student and the process of learning of the student, the more complex model (the teacher) extracted deep level of details from the TCR and peptide structures, then transferred it to a smaller and simpler model (the student). The teacher model contained 64, 128, and 256 filters sequentially applied for three convolutional layers, while the student model contained 16, 32, and 64 filters, respectively. Given these architectures, the student model could reduce the overfit of the teacher model should it happen. Thus, the KD model was used to enhance the generalization capacity of our epiTCR-KDA.

### epiTCR-KDA outperformed existing tools in predicting the binding of unseen peptides

To compare the performance of epiTCR-KDA with currently available tools, we chose a set of benchmarked predictors covering a wide range of data representation and learning approaches, including BERTrand[15], TEIM-Seq[38], TEINet[39], ImRex[8], epiTCR[3] and NetTCR[9], all of which use the CDR3β and peptide sequences as inputs. To ensure a fair benchmark, all participating tools were retrained using the same training set as epiTCR-KDA did (Fig 1A). All tools were tested on 10 non-overlapping testing sets, randomly sampled from the testing data consisting of 1,428,951 CDR3β-peptide pairs. Each testing set contained 60% seen data (pairs of CDR3β-peptide in which the peptide sequences were also found in the training set) and 40% unseen data (pairs of CDR3β-peptide in which peptide sequences were only found in the testing data). This design enabled us to compare the performance of each tool on both seen and unseen data. A significant drop in performance from seen to unseen data indicates low generability of the model.

Overall, epiTCR-KDA performed the best, achieving an average AUC of 0.96 (Figure 2A, Figure S2A and S3A). The second and the third best-performing tools were epiTCR and NetTCR, with average AUC values of 0.92 and 0.89, respectively (Figure 2A, Figure S2A, and S3A). When evaluating their performance on seen data, epiTCR-KDA, epiTCR, and NetTCR showed comparable results with AUC values of 0.99, 0.95, and 0.94, respectively (Figure 2B, Figure S2B). However, on unseen data, epiTCR-KDA clearly outperformed the others, achieving an average AUC of 0.86, compared to 0.54 and 0.57 from epiTCR and NetTCR, respectively (Figure 2C, Figure S2C). Our epiTCR-KDA showed a modest drop in AUC from seen data to unseen data (from 0.99 to 0.86), while the other tools exhibited significant drops (0.95 to 0.54 in epiTCR and 0.94 to 0.57 in NetTCR), suggesting that epiTCR-KDA generalises well.

**Figure 2.**
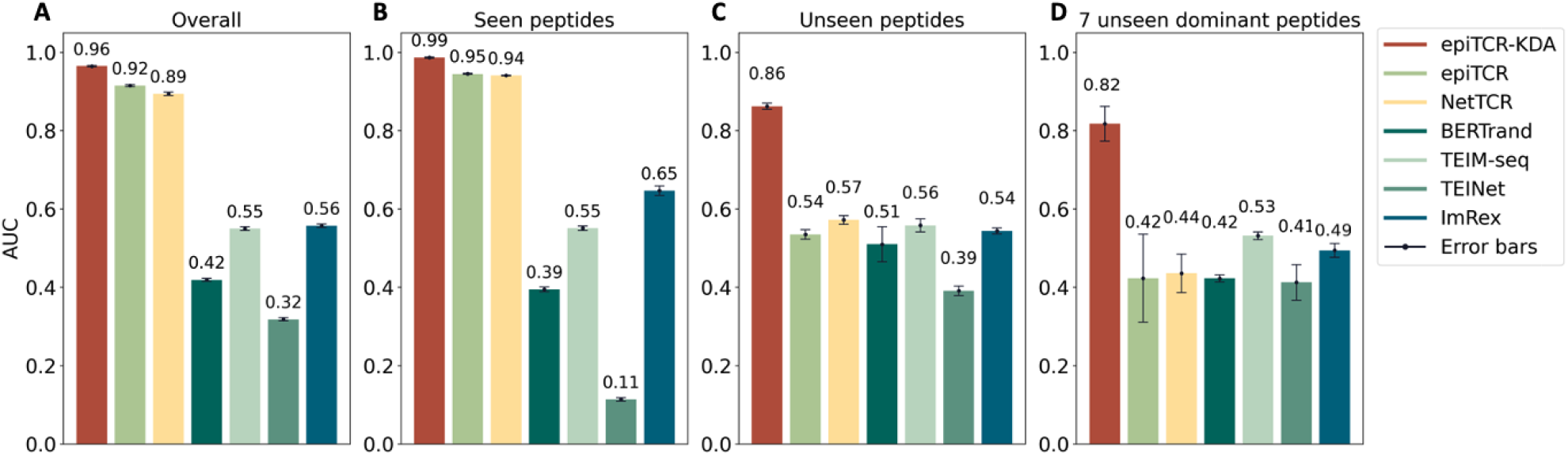
The performance of epiTCR-KDA, epiTCR, NetTCR, BERTrand, TEIM-Seq, TEINet, and ImRex across different benchmark settings: (A) ten overall testing sets including both seen and unseen data, (B) data derived from seen peptides, (C) data derived from unseen peptides, and (D) data derived from 7 dominant unseen peptides (Supplementary Table S3). The performance was measured by AUC. Each bar indicates the mean performance from ten testing sets and the error bar indicates the standard deviation.

To further challenge the models, we included a special testing set of 447,398 CDR3β-peptide pairs derived from 7 unseen peptides, which hereafter referred to as the “7 unseen dominant peptides”[3, 9]. These dominant peptides have been previously observed (Supplementary Table S3), and here we reported the performance of each tool on the CDR3β-peptide pairs derived from these peptides. For all tools tested, the performances on the dominant peptides were slightly lower than those on unseen data (Figure 2D, Figure S2D, and S3D), confirming that data derived from the 7 unseen dominant peptides are more challenging to predict. Despite that, our epiTCR-KDA still maintained a good performance with AUC of 0.82.

### Dihedrals played a pivotal role in maintaining consistently good performance of our epiTCR-KDA

We aimed to understand the factors contributing to the consistent performance of epiTCR-KDA on both seen and unseen data. To achieve this, we evaluated the influence of training data on prediction outcomes, specifically focusing on the similarity between the TCRs and peptides in the training data versus those being predicted. We grouped each testing set into nine clusters based on their CDR3β dihedral angles. Nine representatives were used to represent the diversity of CDR3β sequences across the ten testing sets. For each CDR3β representative, we split the training data into bins containing the CDR3β-peptide pairs, of which the respective CDR3β sequences maintained similar (i.e. in same range of cosine similarity of the phi-psi vectors) to the representative CDR3β (see Supplementary Methods). Next, we calculated the root mean squared error (RMSE) to measure the discrepancy between the labels (binding/non-binding) of the testing cluster and those of the corresponding training bin. We observed a reduction in RSME as the similarity increased across all nine tested CDR3β sequences. It was shown that the binding/non-binding CDR3β-peptide pairs predicted by the epiTCR-KDA in each testing cluster were more associated with those of the training bins exhibiting higher cosine similarity (Figure 3A). We performed similar measurements for the peptides and observed similar patterns (Figure 3B). Overall, these findings suggested that dihedral angles of both CDR3β and peptide could be the key features that determined the outstanding performances of the epiTCR-KDA.

**Figure 3.**
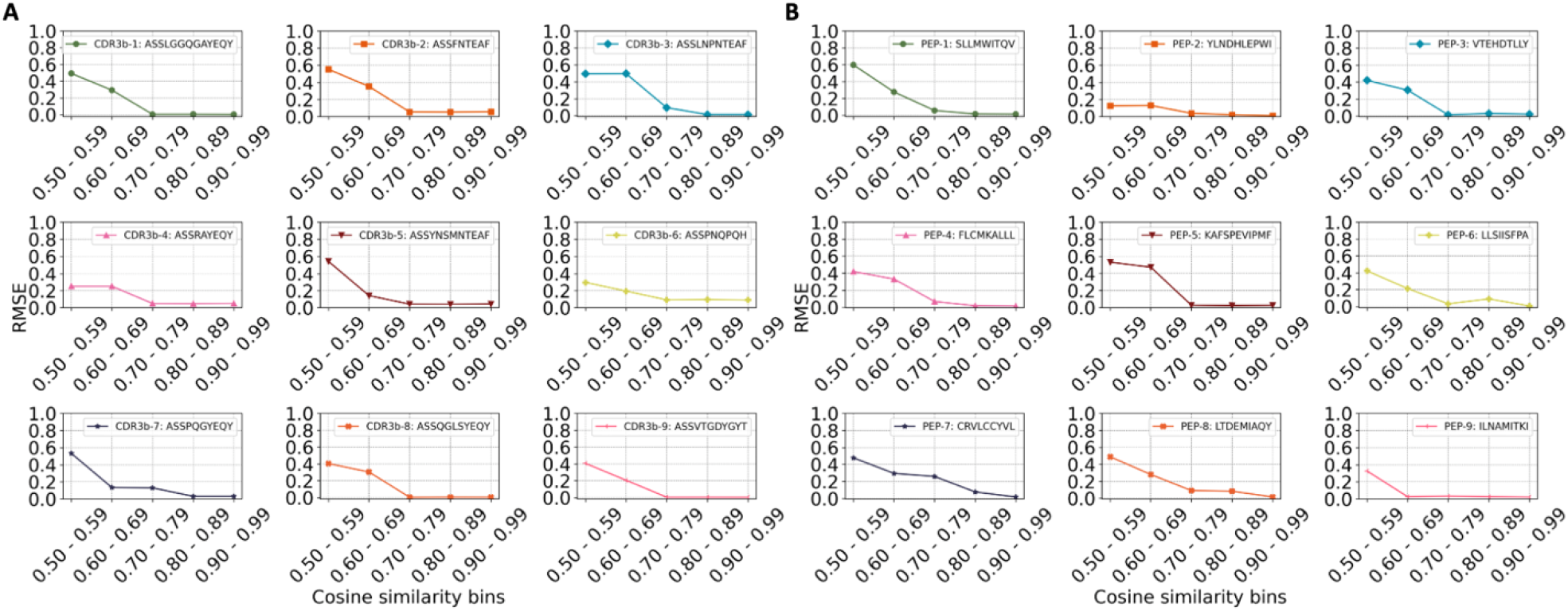
The influence of CDR3β and peptide structural information in training data on predictions by epiTCR-KDA. (A) Nine CDR3β and (B) nine peptides were chosen to represent nine clusters within the testing sets, and the predicted labels of their represented clusters were compared with the labels in training bins at different levels of dihedral angle-based cosine similarity using RMSE. The lower the RMSE, the more similar between prediction labels and training labels.

### Robust performance of the epiTCR-KDA across different testing scenarios

A challenge in the TCR-peptide binding prediction is the generalizability of the prediction models, which might be varied with respect to different testing sets. We demonstrated the potential generalizability and robustness of our epiTCR-KDA by testing its performance across various datasets from multiple sources. The testing data were specifically designed to encompass diverse sources and varying ratios of non-binding to binding CDR3β-peptide pairs. The binding pairs were sourced from two studies: Panpep[40], with 10,397 CDR3β-peptide pairs, and catELMo[38] with 85,020 CDR3β-peptide pairs. The non-binding pairs were generated as previously described (Figure 1A) by combining public CDR3β sequences ([24, 25, 26, 27, 28]) with wildtype peptides([29, 30, 31]). The resulting non-binding sets (n = 402,732 and n = 313,161, Figure 1A) contained no CDR3β-peptide pairs that were present in those used in the earlier benchmark (Figure 2). These binding and non-binding CDR3β-peptide pairs were then combined to form nine new testing datasets (Supplementary Table S4). The performance (AUC) of epiTCR-KDA and six other predictors is shown in Figure 4A, with epiTCR-KDA exhibiting the median AUC of 0.9 (ranging from 0.82 to 0.91), followed by epiTCR achieving the median AUC of 0.88 (ranging from 0.76 to 0.89), and NetTCR reaching the median AUC of 0.79 (ranging from 0.7 to 0.81). It was observed that across the nine testing sets, different performances of the epiTCR-KDA were most likely attributed to the different ratios of unseen-to-seen data.

**Figure 4.**
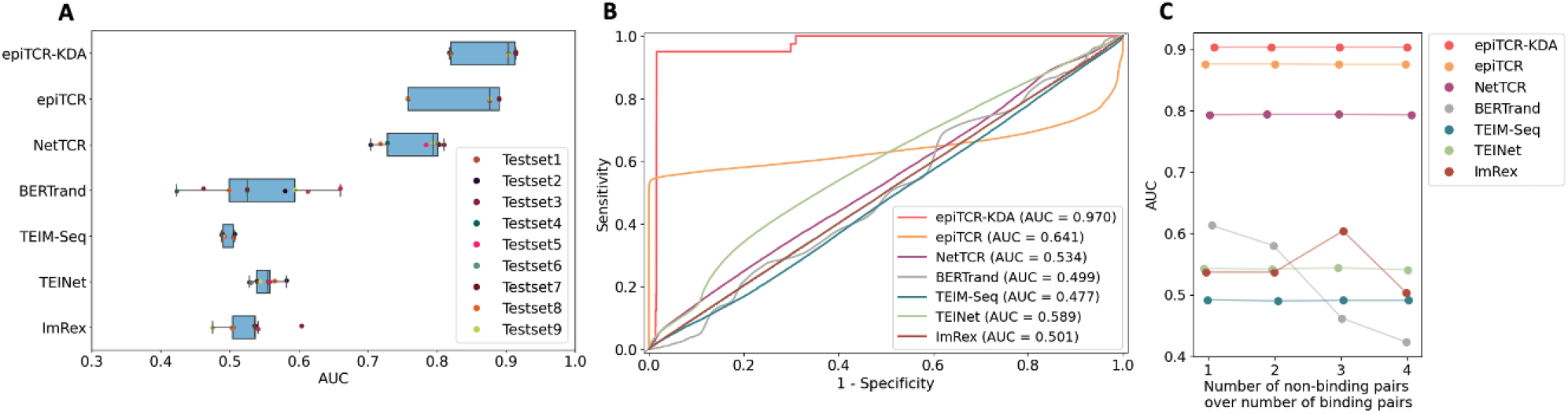
The AUC of epiTCR-KDA, epiTCR, NetTCR, BERTrand, TEIM-Seq, TEINet, and ImRex tested on: (A) nine combined datasets, (B) on the COVID-19 dataset, and (C) on four datasets with an increasing number of non-binding pairs epiTCR-KDA outperformed other TCR-peptide binding predictors with good generalization, while TEIM-Seq, BERTrand, and ImRex remained close to random guesses.

A COVID dataset[41], consisting of 2,120,140 CDR3β-peptide pairs (including 2,120,100 non-binding pairs, and 40 binding pairs), was also used in our subsequent benchmark (Figure 4B). This dataset includes SARS-CoV-2 peptide sequences that were not used in the training data by the epiTCR-KDA. The ratios of seen vs unseen were found in the peptides 1:125.5 and in the CDR3β sequences 1:37.7 (Supplementary Table S4). Despite the more predominant unseen data in this COVID dataset, the epiTCR-KDA performed consistently well (AUC = 0.97) in Figure 4B. The next two best-performing models epiTCR and NetTCR experienced a significant drop in performance (AUC to 0.641 and 0.534, respectively). This result demonstrated the epiTCR-KDA generalizability given the unseen COVID data.

Subsequently, we assessed the performance of epiTCR-KDA and the other predictors using different ratios of the non-binding pairs versus the binding pairs (Figure 4C, Supplementary Table S5). Generally, all the predictors (except for ImRex) performed best when this ratio was 1:1. (Figure 4C). Interestingly, the top three predictors, epiTCR-KDA, epiTCR and NetTCR, consistently performed well even given the increasing ratios of the non-binding versus binding. It suggests that the epiTCR-KDA maintains its robustness.

## Discussion

The potential of using neoantigens as personalised, cancer-specific markers for various therapeutic and preventative anti-cancer strategies has not been fully realised. This is partly due to the difficulties in identifying neoantigens individually for each patient. Numerous computational methods have been developed, employing a wide range of advanced deep learning models to predict TCR-peptide binding (NetTCR[9], TEIM-Seq[42], TEINet[39], BERTrand[15], and ImRex[8]). However, these methods typically rely on amino acid sequences as input, or attempt to convert those sequences using canonical encoding techniques, such as BLOSUM[3, 9], one-hot[39], and physicochemical properties[43]. In this study, we proposed the dihedral angles, also known as Ramachandran angles[44], as input features to predict the TCR-peptide binding (Figure 1). This approach is efficient and captures the three-dimensional structure of both the TCRs and peptides. Although the concept of dihedral angles is well established, to the best of our knowledge its application to predict the TCR-peptide binding pairs has not been reported previously. By providing the angular orientations of consecutive peptide bonds, we hypothesize that our model, epiTCR-KDA, could effectively learn spatial information from CDR3β and peptide, which is crucial for differentiating non-binding from binding CDR3β-peptide pairs. In fact, our epiTCR-KDA model performed consistently well across all testing scenarios (Figure 2 and Figure 4). This was particularly evident in cases where the number of unseen peptides far exceeded the seen peptides, as demonstrated in the COVID dataset (Figure 4B). Our epiTCR-KDA also exhibited high generalizability.

Knowledge distillation has proven effective in various domains, such as natural language processing[45, 46, 47, 48], computer vision[41, 49, 50, 51], and speech recognition[52, 53, 54, 55]. Its versatility stems from its capacity to distill the rich knowledge captured by a complex model into a more compact representation, which is suitable for deployment in environments with limited resources. In the prediction of CDR3β-peptide binding, where accurate modelling of complex molecular interactions is essential, knowledge distillation offers a pathway to enhance the performance of simpler predictive models. By incorporating knowledge distillation with dihedral angles, our model learns from both CDR3β and peptide representations, enabling it to capture a broader range of structural features that influence binding interactions, e.g our epiTCR-KDA exhibited substantial association of both the CDR3β and peptide similarity between training and testing data (Figure 3). In contrast, our previous model, epiTCR, demonstrated that only 50% of the examined peptides had predicted labels similar to those of their corresponding groups of similar peptides in the training set[3]. This finding affirms why epiTCR is less effective than epiTCR-KDA in predicting outcomes on unseen data (Figure 2). Although we have not been able to determine whether this generalization capability is attributable to the dihedral angles input, the KD model, or a combination of both, our findings demonstrate that the epiTCR-KDA represents a promising and novel approach in the area of TCR-peptide binding prediction that has not been previously reported.

Nonetheless, several limitations remain in our current study. First, our data representation is dependent on the reliability of the OmegaFold tool[19] to predict the 3D structures of CDR3β and peptides. We however have not confirmed these resulting 3D models using some other tools such as AlphaFold[18] and ESMFold[19] due to a few scenarios: (1) the short length of the CDR3β and peptide sequences do not satisfy the constraints by AlphaFold (≥16 residues), (2) ESMFold consistently fails for certain of our sequences. Neither did we apply RosettaFold[56] due to our time and resource constraints. Secondly, our model search may not be exhaustive, hence the knowledge distillation model may not be the best model to fully capture the intricacies. Thirdly, while our work has clearly demonstrated that incorporating 3D structure information in data representation can improve the generalizability of our model, there still remain other structural characteristics to be explored to devise more robust and versatile prediction models for diverse CDR3β-peptide complexes. In brief, a model with enhanced interpretability will further advance our understanding of immune system dynamics and facilitate the development of novel therapeutic strategies.

## Conclusion

We presented epiTCR-KDA, a knowledge distillation model that uses dihedral angles for prediction of TCR-peptide binding. By capturing the structural information of both the partners of the TCR-peptide complexes, epiTCR-KDA elicits its generalizability and robustness across diverse datasets. Given its generalizability, the epiTCR-KDA might pave the ways for future development in the areas of immunotherapy that faces low success rate of identifying multiple personalised neoantigens capable of activating T cells.

