## Supplementary File for "epiTCR-KDA: Knowledge Distillation model on Dihedral Angles for TCR-peptide prediction"

### Supplementary Methods

#### Data collection, data preprocessing, and data generation

We presented a comprehensive analysis of CDR3β-peptide interactions by leveraging diverse data sources to gain a broad understanding of the learned model for CDR3β-peptide interactions.

First, we collected binding and non-binding CDR3β-peptide pairs from McPAS-TCR[1], TBAdb[2], VDJdb[3], IEDB[4], and 10X[5]. Each dataset underwent individual preprocessing. For CDR3β, we used amino acid sequences, and where applicable (in TBAdb, VDJdb, McPAS-TCR, and 10X datasets), we removed the "C" starting and "FW" ending characters in the CDR3β chain, similar to NetTCR[6]. In IEDB, we used curated CDR3β sequences. We also eliminated sequences containing unknown amino acids (encoded by X, O, special characters, and lowercase letters). Additionally, we removed empty peptides and unknown amino acids. We filtered all sequences based on their length, with CDR3β lengths ranging from 8 to 19 amino acids and peptide lengths from 8 to 11 amino acids. The combined dataset from these sources comprises 70,083 binding CDR3β-peptide pairs and 2,689,709 non-binding CDR3β-peptide pairs. The 10X dataset included both binding and non-binding samples, with a very small proportion of binding pairs, accounting for less than 1% of the total data.

To create additional non-binding data to increase the diversity of training set, we proposed a strategy for data generation which can be done via two approaches. The first approach involved extracting representative sequences from three publicly available neoantigen datasets—namely, TSNAdb, Neodb, and NEPdb—and randomly combining them with CDR3β sequences from previously collected data (from McPAS-TCR, TBAdb, VDJdb, IEDB, and 10X). This resulted in 174,944 CDR3β-peptide pairs with 2,506 unique peptides in the non-binding dataset. In the second approach, we used 71 random CDR3β sequences from tumor infiltrating lymphocytes (TIL)[7] coupled with peptides from public wild-type datasets (McPAS-TCR, TBAdb, VDJdb, IEDB, and 10X), resulting in 132,979 CDR3β-peptide pairs. These data generation strategies were employed throughout the entire manuscript, including further model evaluations to ensure the non-binding label of the synthesized data.

#### Data organization for model training and testing

The training data affects the learning of the models. Therefore, we first focused on the amount of data needed for model training. From the total 3,647 unique peptides related to interactions gathered in our dataset, we tried to find out the appropriate amount of training data to stabilise the model's performance. Our previous work showed that the model's performance relied significantly on the number of unique peptides. Consequently, we chose an increasing number of peptides and their corresponding interactions for training, specifically 1,000, 1,200, 1,400, 1,600, 1,800, 2,000, 2,400, 2,600, 2,800, 3,000, and 3,200, then performed a 10-fold cross validation training strategy to find the best amount of training data needed for the model generalization. The training peptides were selected based on their frequency (the number of interactions where they appeared), as we prioritized learning from interactions that are common in real life. By gradually increasing the number of peptides, we aimed to strike a balance between data complexity and model generalization. We assessed the model's robustness using the mean AUC across 10 validation folds.

#### Model training

The epiTCR-KDA structure began with the transformation of input CDR3β and peptide sequences into matrices of phi and psi angles. These matrices were then concatenated and zero-padded to form a 17x4 matrix. In this matrix, 17 rows represented the dimension obtained from the longest sequence, and 4 columns represented two pairs of phi and psi angles from both the CDR3β and peptide.

Using this matrix as input, both the student and teacher models were built based on convolutional neural networks (CNNs). The teacher model was designed for binary classification. It started with a convolutional layer of 64 filters of size 3x3 (with a stride of (2, 2)), followed by a LeakyReLU activation ($\alpha = 0.2$), and a MaxPooling2D layer with a 2x2 filter and stride = 1. Subsequently, two convolutional layers with 128 and 256 filters (using the same filter size and stride) were applied. The output from the last layer was flattened into a 1D vector, followed by a fully connected layer and a single unit equipped with sigmoid activation for binary classification.

The student model replicated the teacher's predictions with reduced complexity by reusing three convolutional layers with 16, 32, and 64 filters, respectively, while keeping other layers unchanged from the teacher model. The distillation process involved a Distiller object containing both models. During training, the Distiller object was compiled using Adam optimizer, with BinaryAccuracy metric for evaluation, BinaryCrossentropy loss function for the student, and KLDivergence for distillation loss evaluation. The 'alpha' was set at 0.1 to balance the hard student predictions and the soft distillation loss from the teacher. The 'temperature' was adjusted to 10 to moderate the teacher's predicted probabilities before transferring them to the student. The chosen batch size, set at 64, determined the quantity of training samples processed in each iteration, potentially increasing the training speed and model convergence. These parameters were crucial for achieving effective model performance during training.

#### General model evaluation on remaining data from training

Our initial evaluation of the model's performance focused on the remaining data that had not been utilised during model training. We assessed the model's performance using the Area Under the Curve (AUC) across four scenarios:

1. All CDR3β-peptide pairs: all interactions had not appeared during the training process.
2. Interactions of seen peptides: interactions involving peptides it had encountered during training.
3. Interactions of unseen peptides: interactions with peptides it had not seen during training.
4. Interactions of seven dominant unseen peptides: interactions involving seven specific unseen peptides that were prevalent in the entire dataset.

The full evaluation result is shown in Figure 2, S2, S3, and S4.

#### Evaluating the impact of training data on model's prediction

The process of learning from CDR3β and peptide sequences significantly influenced the model's predictions. To do this, we explored whether the similarity between training and testing CDR3β/peptide sequences affected the model's predictions. Therefore, we compared the predicted interactions of CDR3β/peptides in the testing set with learned interaction labels of similar CDR3β/peptides.

The method thereby described was applied for peptides, and the same calculations were also applied for CDR3β.

Supposing the phi and psi angles of any two peptides were:

$$peptide_i = \{\phi_i, \psi_i\} = \{\phi_{i1}, \phi_{i2}, \dots, \phi_{in-2}, \psi_{i1}, \psi_{i2}, \dots, \psi_{in-2}\} = \{x_1, x_2, \dots, x_{2n-4}\}$$

$$peptide_j = \{\phi_j, \psi_j\} = \{\phi_{j1}, \phi_{j2}, \dots, \phi_{jn-2}, \psi_{j1}, \psi_{j2}, \dots, \psi_{jn-2}\} = \{y_1, y_2, \dots, y_{2n-4}\},$$

with n is the length of peptide sequences.

The similarity between any two peptides relied on the cosine similarity between the phi and psi angles of those peptides. The cosine similarity was defined as:

$$cosine\_similarity_{(peptide_i, peptide_j)} = \frac{\sum_1^{2n-4} x_i y_j}{\sqrt{\sum_1^{2n-4} x_i^2}\sqrt{\sum_1^{2n-4} y_j^2}} \quad (1)$$

given that -1 ≤ $cosine\_similarity(peptide_i, peptide_j)$ ≤ 1.

$$RMSE = \sqrt{\frac{(\%_{pos\_test} - \%_{pos\_train})^2 + (\%_{neg\_test} - \%_{neg\_train})^2}{2}} \quad (2)$$

We selected nine representative CDR3β and nine peptides, each representing distinct groups of CDR3β and peptides available in the testing sets. These groups were formed based on the cosine similarity between CDR3β dihedral angles and/or peptide dihedral angles (equation (1)). For each representative CDR3β/peptide, we organized trained CDR3β/peptides into bins based on their cosine similarity to the representative. The number of CDR3β/peptides having cosine similarity below 0.5 was too small, so we only reported the group of CDR3β/peptides having cosine similarity from 0.5 and above. We categorized training CDR3β/peptides into five levels of similarities: (0.5 – 0.59), (0.6 – 0.69), (0.7 – 0.79), (0.8 – 0.89), and (0.9 – 0.99). We then calculated the Root Mean Square Error (RMSE) reflecting the difference between the labels (binding/non-binding) of CDR3β-peptide pairs related to CDR3β/peptides in the bins and the model's prediction on interactions of CDR3β/peptides under consideration (equation (2)).

#### epiTCR-KDA performance on different testing scenarios

To assess the model's generalizability, we conducted benchmark evaluations on distinct datasets sourced from other research papers. This evaluation occurred in two scenarios.

First, we gathered data from Panpep[8] and catELMo[9]. These datasets exclusively contained binding pairs, necessitating the generation of non-binding pairs using the data generation strategies previously employed for the training and testing sets of our models. The first non-binding dataset comprised 313,161 CDR3β-peptide pairs, consisting of CDR3β sequences from TIL[10] and public wild-type peptides from TSNAdb[11], Neodb[12], and NEPdb[13] (labeled as (1)). The second non-binding dataset included 402,732 pairs from wild-type peptides with public CDR3β sequences (labeled as (2)). Additionally, we utilised binding data from Panpep (10,397 pairs, labeled as (3)) and catELMo (85,020 pairs, labeled as (4)). By exhaustively combining these datasets, we generated a total of nine testing sets (as detailed in Supplementary Table S3). This approach ensured that model evaluation remained independent of any single data source.

In the second scenario, we collected a COVID-19 dataset [14], which consisted of 2,120,140 CDR3β-peptide pairs. Among these, there were 2,120,100 non-binding pairs and only 40 binding CDR3β-peptide pairs. We employed this dataset for an independent testing because the peptides and all related TCRs originated from a context vastly different from the pairs that had been trained and tested by our models.

In the subsequent experiment, we investigated whether the models' performance was affected by the data composition, particularly when we altered the number of non-binding pairs and the amount of unseen peptide interactions. Using the two binding datasets collected from Panpep and catELMo, along with the two non-binding sets generated by our strategies, we adjusted the ratios of binding to non-binding data and the ratios of seen to unseen peptide interactions. Specifically, we created four testing sets with the number of non-binding pairs equal to, double, triple, and quadruple the number of binding CDR3β-peptide pairs. Additionally, we constructed four other CDR3β-peptide sets, each containing the number of unseen peptides equal to five, ten, and twenty times the number of peptides seen in the training set (as detailed in Supplementary Table S4). These data settings were designed to reflect real-life scenarios, where unseen peptide interactions and non-binding interactions significantly contribute to the prediction set.

For all above experiments, the Area Under the Curve (AUC) was reported for the tools participating in the evaluation.

### Supplementary Tables

Table S1. The number of observation in five datasets.

| Datasets | Data counts | Date of collection | Link to dataset |
|---|---|---|---|
| TBAdb | 1,015 | June 16th, 2022 | https://gitlab.com/immunomind/immunarch/blob/master/private/TBAdb.xlsx |
| VDJdb | 7,581 | June 16th, 2022 | https://vdjdb.cdr3.net/search |
| IEDB | 51,680 | June 16th, 2022 | https://www.iedb.org |
| McPAS-TCR | 3,646 | August 5th, 2022 | http://friedmanlab.weizmann.ac.il/McPAS-TCR |
| 10X | 2,689,720 | June 20th, 2022 | 1) https://www.10xgenomics.com/resources/datasets/cd-8-plus-t-cells-of-healthy-donor-1-1-standard-3-0-2<br>2) https://www.10xgenomics.com/resources/datasets/cd-8-plus-t-cells-of-healthy-donor-2-1-standard-3-0-2<br>3) https://www.10xgenomics.com/resources/datasets/cd-8-plus-t-cells-of-healthy-donor-3-1-standard-3-0-2<br>4) https://www.10xgenomics.com/resources/datasets/cd-8-plus-t-cells-of-healthy-donor-4-1-standard-3-0-2 |
| Total | 2,753,642 | | |

Table S2. A table detailing the specific statistics of the collected data and the data generated for the training and testing sets.

| N. | Number of … | | in training set | in testing set |
|---|---|---|---|---|
| 1 | CDR3β-peptide pairs | Binding pairs | 34380 | 29553 |
| | | Non-binding pairs | 1598234 | 1399398 |
| | | Total | 1632614 | 1428951 |
| 2 | Collected CDR3β-peptide pairs | Binding pairs | 34380 | 29553 |
| | | Non-binding pairs | 1463832 | 1225877 |
| | | Total | 1498212 | 1255430 |
| 3 | Generated CDR3β-peptide pairs (non-binding only) | | 134402 | 173521 |
| 4 | Unique peptides | Unique seen peptides | NA | 1948 |
| | | Unique unseen peptides | NA | 1641 |
| | | Collected unique peptides | 159 | 1174 |
| | | Generated unique peptides | 1894 | 2444 |
| | | Total | 2,000 | 3589 |
| 5 | Unique CDR3β | Unique seen CDR3β | NA | 63237 |
| | | Unique unseen CDR3β | NA | 26613 |
| | | Collected unique CDR3β | 87375 | 89828 |
| | | Generated unique CDR3β | 71 | 71 |
| | | Total | 94725 | 89850 |

Table S3. List number of pairs derived from each of the 7 dominant peptides

| No | Peptide | n_pairs |
|---|---|---|
| 1 | GLCTLVAML | 65,713 |
| 2 | NLVPMVATV | 65,109 |
| 3 | GILGFVFTL | 64,111 |
| 4 | TPRVTGGGAM | 63,959 |
| 5 | ELAGIGILTV | 63,495 |
| 6 | AVFDRKSDAK | 62,774 |
| 7 | KLGGALQAK | 62,237 |

Table S4. Counts of CDR3β-peptide pairs in different testing sets used in Figure 4A and 4B

| No | Dataset | n_obs | n_pos_obs | n_neg_obs | n_unique_pep | n_unseen_pep | n_seen_pep | n_unique_tcr | tcr_unseen | tcr_seen |
|---|---|---|---|---|---|---|---|---|---|---|
| 1 | (1)(3) | 398181 | 85020 | 313161 | 3540 | 1756 | 1784 | 81780 | 81777 | 3 |
| 2 | (1)(4) | 323558 | 10397 | 313161 | 3014 | 1286 | 1728 | 10416 | 10416 | 0 |
| 3 | (2)(3) | 487752 | 85020 | 402732 | 3540 | 1756 | 1784 | 81694 | 81691 | 3 |
| 4 | (2)(4) | 413129 | 10397 | 402732 | 3014 | 1286 | 1728 | 10276 | 10276 | 0 |
| 5 | (1)(3)(4) | 408578 | 95417 | 313161 | 3553 | 1764 | 1789 | 86743 | 86740 | 3 |
| 6 | (2)(3)(4) | 498149 | 95417 | 402732 | 3553 | 1764 | 1789 | 86656 | 86653 | 3 |
| 7 | (1)(2)(3) | 800913 | 85020 | 715893 | 3540 | 1756 | 1784 | 81858 | 81855 | 3 |
| 8 | (1)(2)(4) | 726290 | 10397 | 715893 | 3014 | 1286 | 1728 | 10544 | 10544 | 0 |
| 9 | (1)(2)(3)(4) | 811310 | 95417 | 715893 | 3553 | 1764 | 1789 | 86815 | 86812 | 3 |
| 10 | COVID | 2120140 | 40 | 2120100 | 1265 | 1255 | 10 | 1489 | 1442 | 47 |

Note:

(1): non-binding pairs made of CDR3β from TIL[10] and wildtype peptides,
(2): non-binding pairs made of publicly collected CDR3β[1, 2, 3, 4, 5] and wildtype peptides[11, 12, 13],
(3): binding pairs from Panpep[8],
(4): binding pairs from catELMo[9]

Table S5. Counts of CDR3β-peptide pairs in different testing sets used in Figure 4C

| No | Dataset | n_obs | n_pos_obs | n_neg_obs | n_unique_pep | n_unseen_pep | n_seen_pep | n_unique_tcr | n_unseen_tcr | n_seen_tcr |
|---|---|---|---|---|---|---|---|---|---|---|
| 1 | n.neg = n.pos | 190834 | 95417 | 95417 | 3553 | 1764 | 1789 | 86815 | 86812 | 3 |
| 2 | n.neg = 2n.pos | 286251 | 95417 | 190834 | 3553 | 1764 | 1789 | 86815 | 86812 | 3 |
| 3 | n.neg = 3n.pos | 381668 | 95417 | 286251 | 3553 | 1764 | 1789 | 86815 | 86812 | 3 |
| 4 | n.neg = 4n.pos | 477085 | 95417 | 381668 | 3553 | 1764 | 1789 | 86815 | 86812 | 3 |

### Supplementary Figures

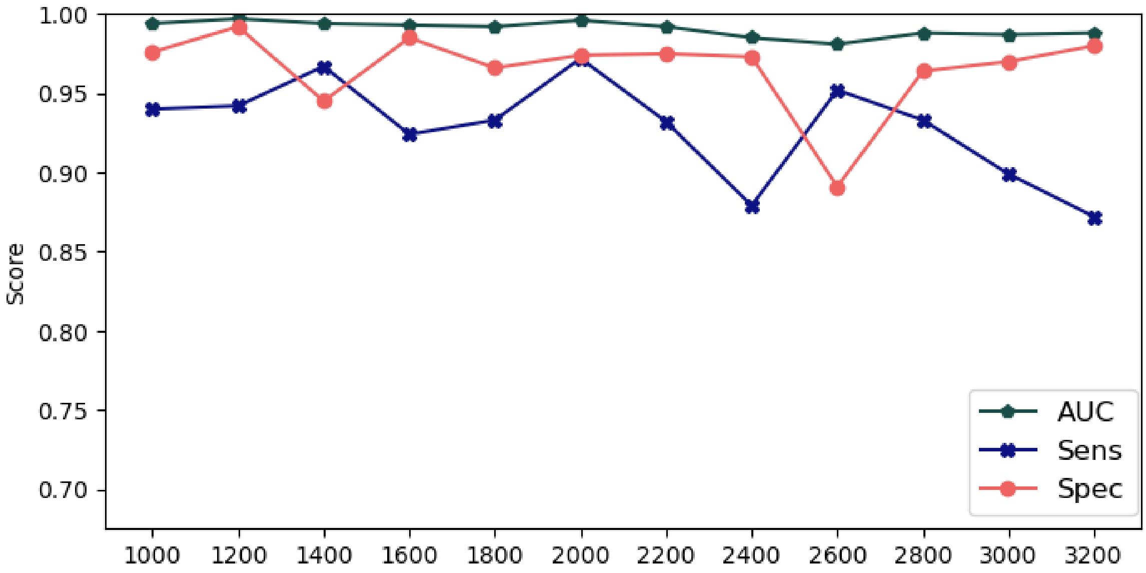


Figure S1. Model performance across different amount of training data, starting from interactions of 1,000 peptides to 3,200 peptides. The reported values were the mean model performance across 10 folds of training validation.

Our testing on gradual changing amount of training data (Figure S1) showed that the top 2,000 peptides led to the good performance in AUC. Furthermore, the model trained with 2,000 peptides had very small trade-off between the model sensitivity and specificity. Therefore, the model trained on this amount of training data was expected to have robust prediction.

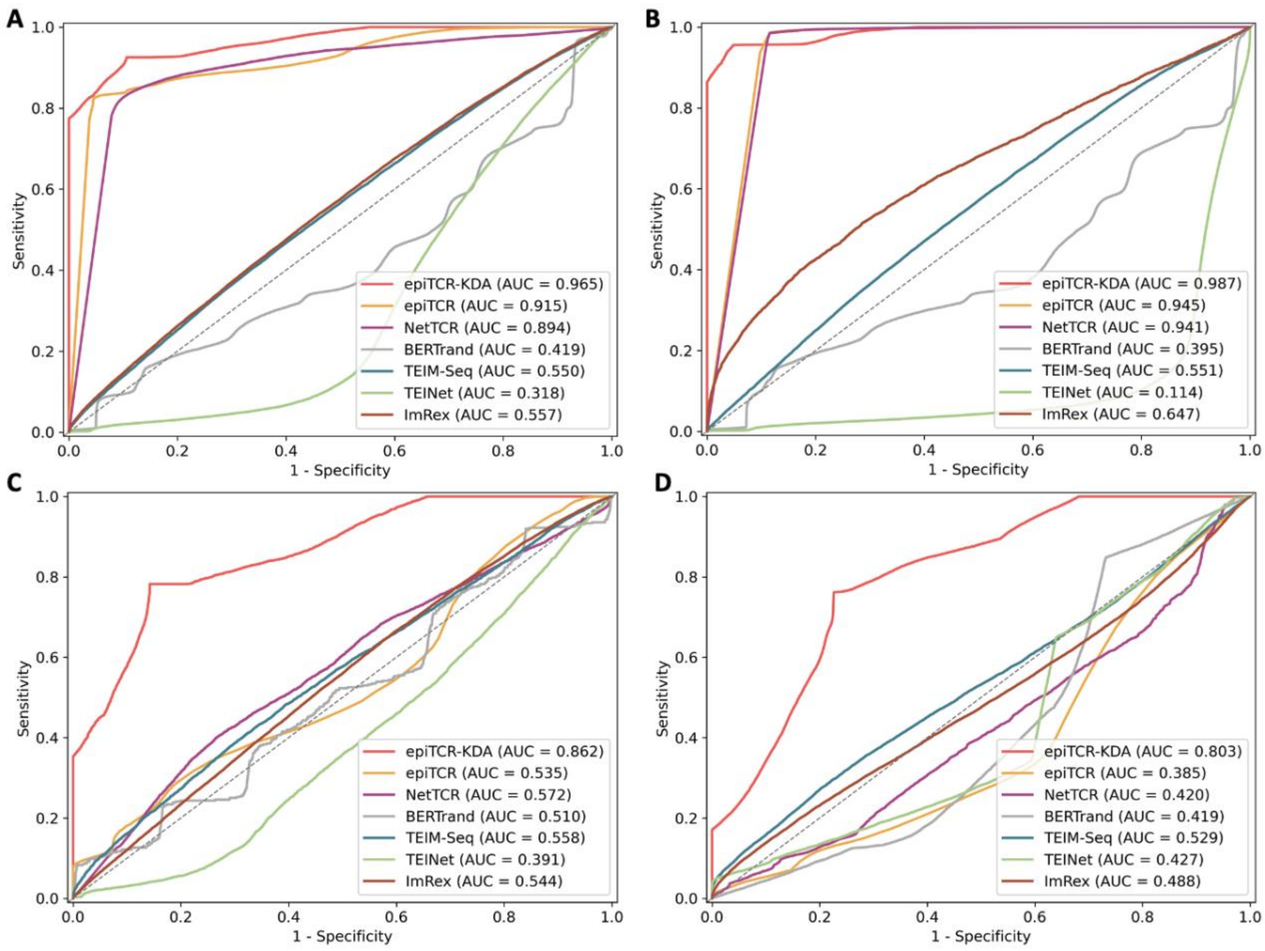


Figure S2. The prediction AUC of epiTCR-KDA, epiTCR, NetTCR, BERTrand , TEIM-Seq, TEINet and ImRex on ten overall testing sets (data remaining from training set), with four benchmark settings: (A) on overall interactions, (B) on interactions of seen peptides, (C) on interactions of unseen peptides, and (D) on interactions of seven dominant unseen peptides.

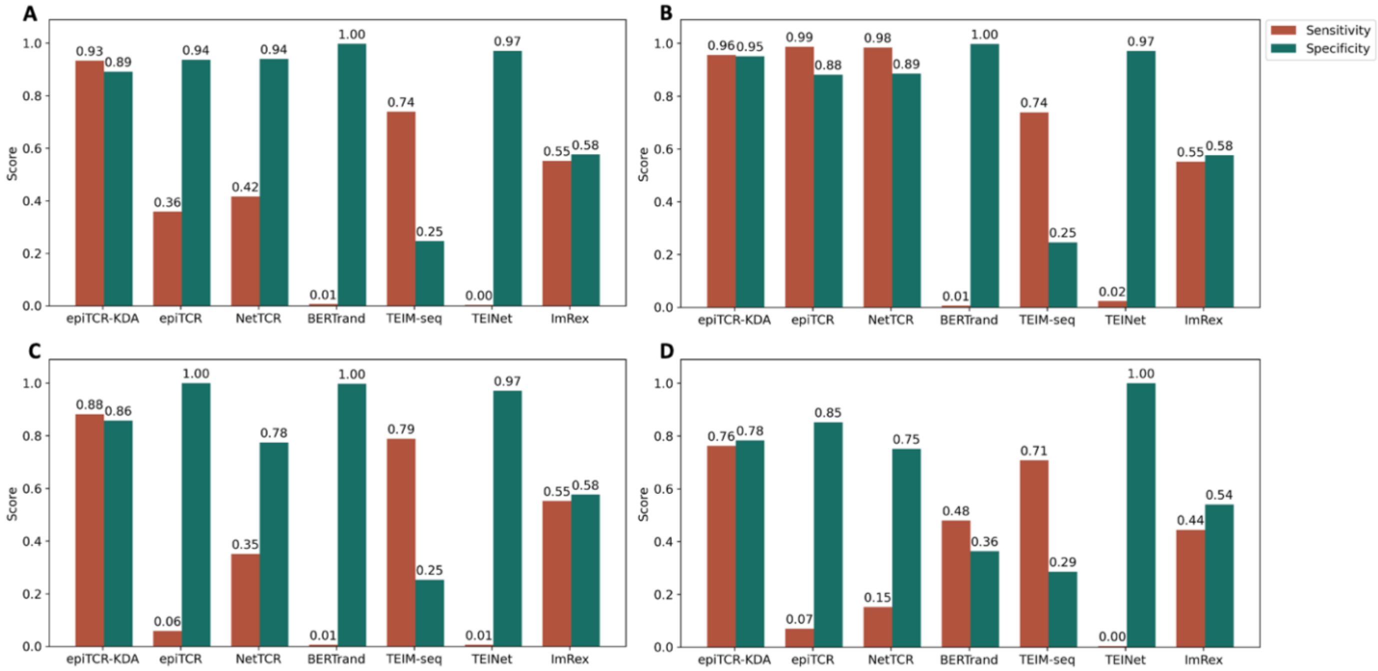


Figure S3. The sensitivity and specificity of the epiTCR-KDA, epiTCR, NetTCR, BERTrand, TEIM-Seq, TEINet and ImRex on ten overall testing sets (data remaining from training set), on four benchmark settings: (A) on overall interactions, (B) on interactions of seen peptides, (C) and on interactions of unseen peptides, and (D) on interactions of seven dominant unseen peptides.
